# Indiv-Brain: Individualized Brain Network Partition Learned from Individual fMRI Data using Deep Clustering with Vertex-level Attention

**DOI:** 10.1101/2024.08.22.608966

**Authors:** Zhitao Guo, Yongjie Duan, Le Xing, Xiaojie Zhao, Li Yao, Zhiying Long

## Abstract

Individualized functional network partitioning is increasingly critical in elucidating individual differences in cognition, development, and behavior. The previous studies grouped brain regions that were pre-defined by common brain parcellations into individualized brain networks. The employment of common brain region parcellations ignores the individual variations in brain structure and reduces the spatial resolution of the results. Moreover, some studies trained models on a group of subjects to guide individual network partitioning. These methods largely depend on the sample size and encounter challenges in the case of limited subject data. In this paper, we propose Indiv-Brain which automatically partition brain vertices into different brain networks based on individual fMRI data without any prior brain parcellation. The Indiv-Brain consists of three sequential modules: stacked denoising autoencoder (SDAE) for mapping the raw fMRI data to a latent embedding space, masked vertex modeling (MVM) for learning attention-enhanced representations of brain vertices, and deep embedding clustering with spatial attention (DEC-A) for unsupervised clustering on the learned vertex representations. The experiments on Human Connectome Project (HCP) demonstrate that the accuracy of Indiv-Brain outperforms existing methods. We compared the model with methods like SDAE, DEC, IDEC, and *k*-means. For the results of 8 subjects, the accuracy of Indiv-Brain was consistently the highest, averaging 6.65 percentage points higher than the IDEC method. To the best of our knowledge, this is the first study to obtain individualized brain network partition based on individual fMRI data with deep learning models. Our study provides a novel insight into understanding individualized brain networks, especially suitable for special patients.

## 1 Introduction

Understanding the functional organization of human brain is a long-standing goal in neuroscience. As a non-invasive imaging method, functional Magnetic Resonance Imaging (fMRI) is one of the most widely used tools via measuring the blood oxygen level dependence (BOLD) signal. Various studies [1–5] demonstrate that human brain can be grouped into several functional networks (subsystems or communities), which coordinate with each other to accomplish complex cognitive processes [6, 7].

However, these findings were based on a group of subjects and ignored the individual variability highlighted by increasing studies [8, 9].Investigating of functional brain variations among individuals is crucial for further understanding individual differences in cognitive processing and may facilitate the discovery of neurological biomarkers of some brain disorders.

Great efforts have been made to explore individualized functional brain networks. These studies can be classified into two principal categories. The first category divides the brain into different regions of interest (ROI) using anatomical or functional brain parcellations to the neuroimaging data and groups the ROIs into different brain network for individuals [10–13]. These studies largely rely on common brain atlases that may not be suitable for some special subjects, and the partitioned brain networks are ROI-based with a coarse spatial resolution. The second category obtains individual-level brain network partition based on group-level algorithms which require a group of subjects [14–19]. However, some of these methods are limited by the assumption that the loadings of corresponding functional networks of different individuals follow certain distributions [15, 17, 18]. Moreover, biased results may be generated if group-level algorithms are applied to individual subjects that are heterogeneous to the group. Therefore, it is essential to investigate individualized brain network partitions without any predefined brain parcellations or other subjects. One of the main advantages of fMRI is its high spatial resolution. For example, there are tens of thousands of vertices/voxels in the surface-style/volume-style fMRI data [20]. Partitioning brain networks at the vertice/voxel level can not only provide more insights into brain modularity at a fine spatial resolution but also eliminate the dependence on any brain atlases or other subjects.

The main challenge of brain network partition at the vertice/voxel level is to cluster the vertices/voxels into networks in an unsupervised learning manner. In recent years, deep-learning-based clustering (deep clustering) algorithms have been proposed and widely applied to unsupervised clustering [21–25]. Deep clustering generally comprises two stages:first, learning the nonlinear mapping from the original data to the hidden feature space via a deep neural network, and second,clustering on the learned latent features. Although deep clustering can be used to cluster the brain vertices/voxels into different brain networks, it can only learn the latent feature representation of each vertices/voxel independently without considering the relationships between vertices/voxels. Due to the spatial correlation of fMRI data, the relationships between vertices/voxels can provide valuable guidance during the clustering stage.

Recently, self-supervised pretraining Transformer models on large-scale data that can learn contextual knowledge from sequences have been successfully applied to natural language processing (NLP) [26] and computer vision (CV) [27–29]. In this study, we proposed Indiv-Brain which incorporates self-supervised pre-training Transformers to capture vertex relationships and guide the latent feature learning of vertices into the deep clustering model.

The tremendous success of self-supervised pretraining on large-scale data in natural language processing (NLP) [26] and computer vision (CV) [27–29] proves that useful context knowledge can be learned. Inspired by this success, we proposed Indiv-Brain. To the best of our knowledge, this is the first study to obtain individualized brain network partition based on individual fMRI data with deep learning models. Our contributions are summarized as follows:

- **Novelty**: We proposed Indiv-Brain that can automatically partition the whole brain vertices into different brain networks based on individual resting-state fMRI data. Indiv-Brain consists of stacked denoising autoencoder (SDAE), masked vertex modeling (MVM) and deep embedding clustering with vertex-level attention (DEC-A).
- **Flexibility**: The proposed model can be applied to any subject without any prior information. Moreover, the model can accommodate various neuroimaging data, including volume or surface-based data.
- **High accuracy**: The accuracy of Indiv-Brain was consistently the highest, averaging 6.65 percentage points higher than the IDEC method.
- **Interpretability**: The visualization of the clustering results and the analysis of the attention mechanism suggest that our model is interpretable in neuroscience.

## 2 Related Work

### 2.1 Representation learning based on autoencoders

Representation learning is a powerful technique that aims to extract meaningful features from raw data. Autoencoder [30], which consists of an encoder mapping the input data into a latent space and a decoder reconstructing the input, is a classical model in representation learning. Denoising autoencoder (DAE) [31] is a class of autoencoders which corrupt the input data with noise and attempt to reconstruct the original, uncorrupted data. A specialized form of DAE is the masked autoencoder (MAE),where parts of the input data are masked and the target is to reconstruct the missing parts. This architecture has been successfully applied in NLP [26], CV [27, 32, 29]and neuroscience [33, 34].

In NLP, masked language modeling (MLM) was proposed in BERT [26], aiming to predict the randomly masked words in a sentence. By understanding the contextual relationships between words, BERT achieved remarkable performance across downstream tasks. In CV, masked image modeling (MIM) has been proposed and used [27–29]. For instance, BEiT [32] transforms the image into visual tokens and predicts the masked discrete tokens. MAE [27] reconstructs the input image by predicting the pixel values of each masked image patch. Similar to MLM and MIM, masked brain modeling (MBM) was proposed and applied in neuroimaging data [33, 34]. The authors in [34] divided fMRI voxels into patches and the prediction target is to recover the masked patches with the voxel values.

### 2.2 Deep clustering

The powerful ability of deep learning to extract informative representations from data has led to unprecedented performance in deep learning-based clustering (deep clustering). The deep embedded clustering (DEC) [21] algorithm performs clustering on the features derived from a nonlinear mapping (SDAE) and then jointly optimizes both the mapping parameters and cluster centers in an unsupervised manner. Improved deep embedded clustering [35] further improves performance by introducing a new loss, which helps to guarantee the local structure preservation of data points. Similarly, [22] proposes a deep clustering network, using the loss function of K-means to help autoencoder learn a “K-means-friendly” data representation. Variational deep embedding [23] proposes an unsupervised generative clustering method that models the data generative procedure with a Gaussian Mixture Model and a deep neural network. DeepCluster[24] proposes an end-to-end method which iteratively clusters deep features learned from CNN and uses the cluster assignments as pseudo-labels to further optimize the parameters.

## 3 Methodology

### 3.1 Overview

The proposed Indiv-Brain model takes the fMRI BOLD signal data ***X*** ∈ ℝ^*n*×*t*^, with *n* vertices and *t* time points, as input and outputs the clustering results of the vertices. As illustrated in Fig. 1, Indiv-Brain comprises three sequential modules: stacked denoising autoencoder (SDAE), masked vertex modeling (MVM), and deep embedding clustering with vertex-level attention (DEC-A). In SDAE, the fMRI time series are transformed into a new latent space through a non-linear mapping ***f***_*θ*_ : ***X***→ ***Z*** ∈ ℝ^*n*×*z*^ via a stacked denoising autoencoder. In MVM, the module learns attention-enhanced representations of all the vertices. Finally, clustering is performed on the learned representations. The design of the framework is flexible, which allows for easy adaptation of new models if better architectures of either one are available.

**Figure 1:**
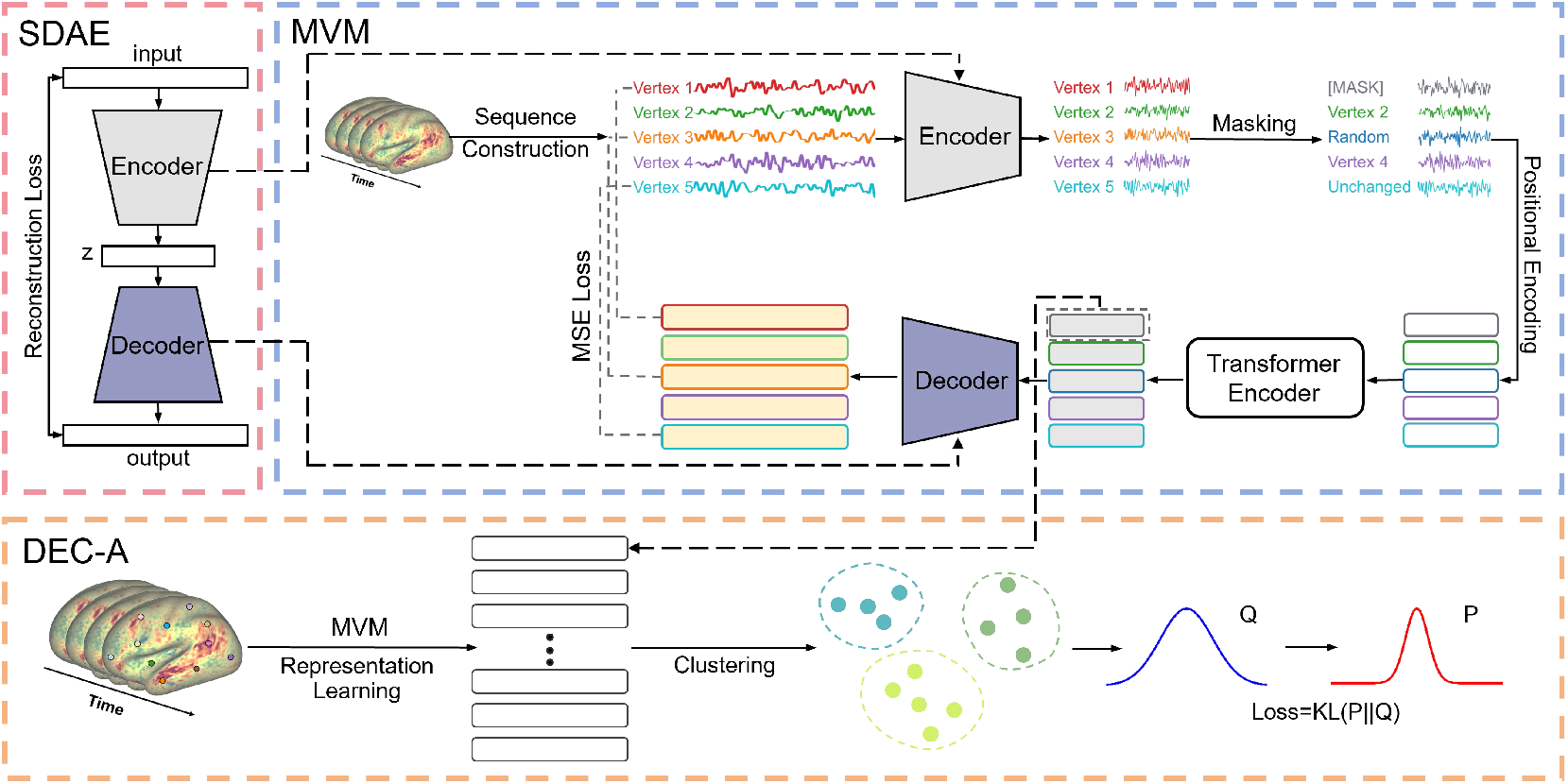
The overview of Indiv-Brain. **Top Left:** Stacked denoising autoencoder (SDAE). **Top Right:** Masked vertex modeling (MVM). **Bottom:**.Deep embedded clustering with vertex-level attention (DEC-A).

### 3.2 Indiv-Brain

#### 3.2.1 Stacked denoising autoencoder (SDAE)

We train the SDAE layer by layer, and each layer is a denoising autoencoder with two-layer MLP defined as:

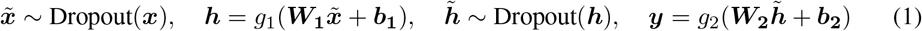

After all the autoencoders are trained, the encoder and decoder of autoencoders are combined to an SDAE and finetuned. Finally, only the encoder part of the SDAE is remained and used as the non-linear mapping in follow-up modules.

#### 3.2.2 Masked vertex modeling (MVM)

##### Sequence construction

Unlike typical NLP tasks where sequences are consist of tokens (words) arranged according to the pre-defined grammar, constructing a sequence in neuroimaging data is challenging since there is no direct semantic relationship between the surface vertices on the brain. Inspired by the “visual grammars” [36] and the widely accepted practices in neuroscience that using Pearson correlation coefficients to measure the strength of functional connections between ROIs, we propose a novel method to build sequences. Specifically, for each vertex, we calculate the correlation coefficients between the time series of the target vertex and all the other vertices. Then we select the top *k*− 1 vertices that are most correlated with the target vertex to form a sequence with length *k* (the first is the target vertex). Finally, for each vertex, we can construct a corresponding sequence starting with it. This method of constructing sequences is data-driven and without the prior knowledge.

##### MVM

While autoencoder is able to learn the informative representations from the time series of each vertex, it ignores the relationship between vertices. In Indiv-Brain, MVM has an encoder-decoder architecture, with the significant difference of incorporating self-attention mechanism to learn the attention-enhanced representations of vertices and the proposed masked vertex modeling (MVM). In MVM, the encoder and decoder are exactly same as those in SDAE. During training, the parameters of encoder and decoder are initialized with the pretrained SDAE.

Given the input BOLD signal data ***X*** ∈ ℝ^*n*×*t*^ with *n* vertices and *t* time points (dimension), the input is transformed to ***X*** ∈ ℝ^*n*×*k*×*t*^ after sequence construction, where *k* is the sequence length. And the data was mapped into a latent space by the encoder: ***X*** ∈ ℝ^*n*×*k*×*t*^ → ***Z*** ∈ ℝ^*n*×*k*×*z*^ via the encoder. Furthermore, the vertex sequences are processed by MVM. Similar to BERT, 15% of the tokens in each sequence are randomly selected. For each selected token, there are three different processing ways: 1)80% probability of being replaced with a learnable parameter [MASK]; 2) 10% probability of being replaced with another vertex; 3) 10% probability of keeping unchanged. We employ a learnable positional encoding ***E*** ∈ ℝ^*k*×*z*^, followed by Transformer encoder, which is based on the original implementation as [37] described. Finally, attention-enhanced sequences are recovered to the original shape ***X*** ∈ ℝ^*n*×*k*×*t*^ through the SDAE decoder. The reconstruction loss function computes the mean squared error (MSE) between the reconstructed and original time series.Note that the objective is to reconstruct the time series of selected vertices instead of all vertices.

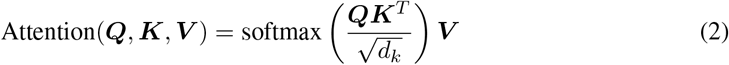

#### 3.2.3 Deep embedded clustering with vertex-level attention (DEC-A)

After extracting the attention-enhanced representations of vertices, DEC-A is employed to perform clustering on the vertices. Deep embedded clustering (DEC) [21] is a two-phase clustering algorithm. Firstly, the raw data is mapped from the data space to a latent space using a non-linear mapping. Then clustering is performed on the *Z* space. A Kullback–Leibler (KL) loss is designed to simultaneously optimize the cluster centers and mapping parameters. The brief description of DEC can be summarized as follows and refer to [21] for more details.

In Indiv-Brain, The MVM is used as the initial mapping ***f***_*θ*_ : ***X*** → ***Z*** that transforms the raw data to latent space. And the initial cluster centroids 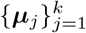 are obtained by employing *k*-means cluster on the data in the space ***Z***. The similarity between embedded points ***z***_*i*_ and centroids ***µ***_*j*_ is calculated by:

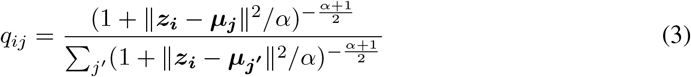

where *α* is set to 1 in our study. Furthermore, target distribution *p*_*ij*_ is calculated based on *q*_*ij*_:

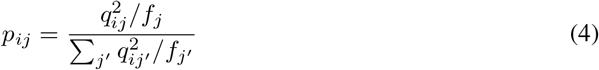

Finally, the *KL* divergence loss between the soft assignments *q*_*ij*_ and the target distribution *p*_*ij*_ is defined as follows:

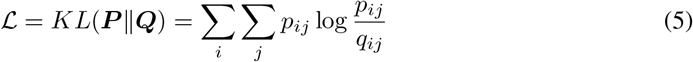

## 4 Experiments

### 4.1 Data preprocessing

We conducted a rigorous preprocessing. See Appendix A for more details.

#### Personalized dataset

Due to individual variability and the absence of a universally accepted “gold standard”, we utilized the widely recognized Yeo atlas [1] as the ground truth. The original Yeo template and the preprocessed fMRI data were in different surface spaces: the FreeSurfer surface space and the 32k_fs_LR surface space, respectively. The latter includes 64,984 vertices, with each hemisphere comprising 32,492 vertices. Vertices within the medial wall were excluded from analysis because this area does not correspond to any defined gray matter structure, resulting in a practical vertex count of 59,412, 29,696 for the left hemisphere and 29,716 for the right hemisphere.

We used a version of the Yeo template that is registered to the 32k_fs_LR space from [38]. It is worth noted that the medial wall of registered Yeo atlas were shifted compared to the standard 32k_fs_LR space, likely due to suboptimal registration/resampling process. Consequently, there is an incomplete match between the non-medial wall vertices of the registered Yeo atlas and those of the standard 32k_fs_LR space. To address this misalignment, we conducted an intersection of the non-medial wall vertices from both surface spaces, resulting in a consistent vertex count of 58,666 for each subject.

### 4.2 Implementation

#### SDAE pretraining

In order to be consistent with previous studies [4, 29], we set the encoder dimension of the SDAE to *d*− 500 −500− 2000 − *z*, where *d* and *z* are the dimensions of input (i.e., 1200) and embedding dimension. We evaluated different embedding dimensions *z* for {10, 32, 64, 128}. During the layer-wise pretraining, each denoising autoencoder is pretrained for 200 epochs with a dropout rate of 20%. During the finetune stage, the entire SDAE is trained for 200 epochs without dropout. The training loss is optimized with the SGD optimizer, the momentum is 0.9 and weight decay is 0. The initialized learning is set to 0.1 and multiplied by 0.1 every 100 epochs. The batch size is 256.

#### MVM pretraining

Similar to the SDAE, we also used MSE loss as the loss function for pretraining. For the embedding dimension, we experimented with the combinations {10, 32, 64, 128}. For each embedding dimension, we obtained the corresponding model parameters from the SDAE model and used them as the initial parameters for the MVM model. Specifically, we employed two different numbers of Transformer encoder blocks {1, 2} and two different sequence lengths {64, 128}. Each training session was run for 1000 epochs. Regarding the learning rate, we chose two peak values {0.0001, 0.00015} and used a warm-up strategy for learning rate adjustment, with the peak values occurring at the 100th epoch and the 1000th epoch, respectively. For the representative subject, the combination of 128 embedding dimensions (hid), 2 blocks (blks), a peak learning rate of 0.00015, and a peak learning rate at the 100th epoch yielded the best results. We used the Adam optimizer for our training. The number of attention heads *n*_*α*_ is determined by 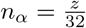, which is consistent with [39]. GELU [40] activation function is used. We randomly select 15% of the tokens (time series of vertices) in every sequence, keeping consistent with BERT [26]. The dropout rate is set to 0.1.

#### DEC-A training

After obtaining the representations of all the vertices, *k*-means is employed to initialize the centroids. Specifically, we run *k*-means for 20 times and select the best solution. During the KL divergence minimization stage, the optimizer is SGD with the momentum of 0.9. The batch size is 256. We trained 50 epochs with the fixed learning rate of 0.01.

#### Computing resource

The experiments were conducted on a high-performance computing setup. The system was equipped with an AMD EPYC 9754 128-Core Processor CPU and an NVIDIA GeForce RTX 3090 GPU with 24 GB of VRAM. The operating system used was Ubuntu 20.04 LTS. Our model is implemented in Pytorch [41]. Training times vary depending on model size but were generally in about 6 hours for MVM pretraining and less than one hour for DEC-A training.

### 4.3 Evaluation Metrics

We employ the popular metric, unsupervised clustering accuracy (ACC), to evaluate the performance of different algorithms. The ACC can be formulated as:

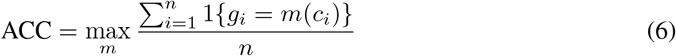

where *g*_*i*_ is the ground truth, *c*_*i*_ is the cluster assignment obtained by the algorithm, and *m* is the map between labels of the ground and the output of algorithms.

## 5 Results

### 5.1 Hyper-parameter evaluation

#### Hyper-parameters of SDAE and DEC/IDEC

We conducted the experiments on the MNIST [42] dataset to verify the reasonableness of the hyperparameter settings.The accuracy and Normalized Mutual Information (NMI) metrics were used to evaluate the performance of our model. The accuracy and NMI of the three methods are: 82.37% and 76.71% for SDAE+*k*-means, 86.96% and 84.69% for DEC, 88.15% and 86.81% for IDEC. The experiment results showed that both the accuracy and NMI scores of the three methods are higher than those reported in the original papers [21, 35], indicating the reasonableness of the hyperparameters settings in pretraining SDAE and training DEC/IDEC.

#### Hyper-parameters of Indiv-Brain

One of the major innovations of our study is that the pretraining is based on individual fMRI data. Therefore, due to the unique nature of our model, there is no single set of hyperparameters applicable to all subjects. We conducted a grid search on the representative subject for four hyperparameters: sequence length, number of attention blocks, embedding dimension, and peak learning rate, resulting in 24 combinations. The accuracy for each combination is shown in Fig. 2. The x-axis represents four hyperparameter combinations: 128len-2blks, 128len-1blk, 64len-2blks, 64len-1blk; the y-axis represents six hyperparameter combinations: 10hid-0.0001lr, 10hid-0.00015lr, 64hid-0.0001lr, 64hid-0.00015lr, 128hid-0.0001lr, 128hid-0.00015lr. For the representative subject, the combination of 128 sequence length, 2 attention blocks, 128 embedding dimension, and 0.00015 peak learning rate achieves the highest accuracy. This set of hyperparameters is not universally applicable to all subjects, where the highest accuracy for each subject may vary across different embedding dimensions. Therefore, our pretrained MVM requires hyperparameter adjustment before being applied to other subjects.

**Figure 2:**
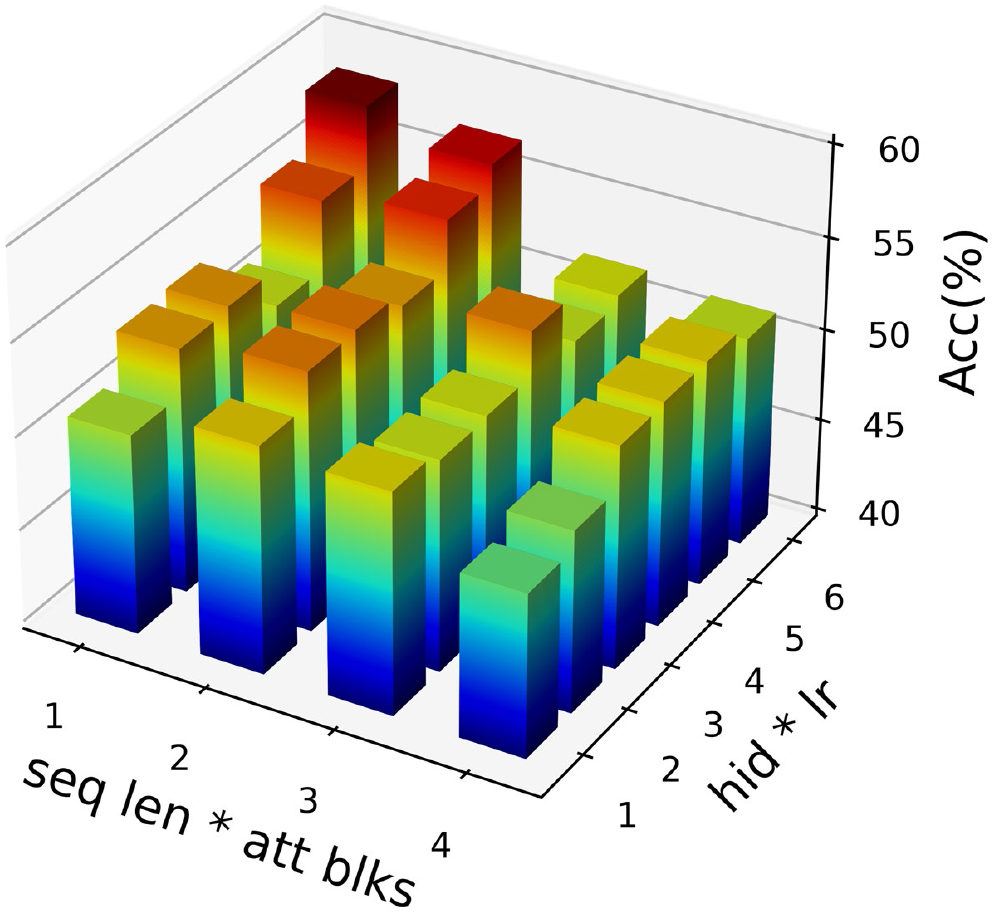
The grid search results for the representative subject. The X-axis and Y-axis correspond to the combinations of 4 hyperparameters, and the Z-axis represents the accuracy obtained compared to the yeo7 template [1].

**Table 1:** The results of the 8 subjects for each model, and values represent the accuracy of the clustering results compared to the yeo7 template [1]. For each subject, the best-performing embedding dimension value was selected for display.

| Subject | 1 | 2 | 3 | 4 | 5 | 6 | 7 | 8 |
| --- | --- | --- | --- | --- | --- | --- | --- | --- |
| hid | 128 | 32 | 32 | 10 | 128 | 64 | 128 | 32 |
| SDAE+*k*-means | 49.49 | 54.18 | 46.92 | 47.75 | 46.52 | 53.93 | 44.97 | 51.23 |
| SDAE+DEC [21] | 50.14 | 54.54 | 48.79 | 48.02 | 46.66 | 53.93 | 45.69 | 52.73 |
| SDAE+IDEC[35] | 50.23 | 55.43 | 50.05 | 48.47 | 48.96 | 53.94 | 41.62 | 54.04 |
| MVM+*k*-means | 54.57 | 61.89 | 53.67 | 46.51 | 49.59 | 57.99 | 45.29 | 52.20 |
| Indiv-Brain | **58.28** | **64.31** | **57.02** | **53.66** | **51.55** | **60.74** | **52.92** | **57.49** |

### 5.2 Interpretation Results

After SDAE pretraining, MVM pretraining, and DEC-A training, each vertex is assigned to a cluster. For each method, we project the vertices onto the fs_LR_32k brain surface, as shown in Fig. 3.

**Figure 3:**
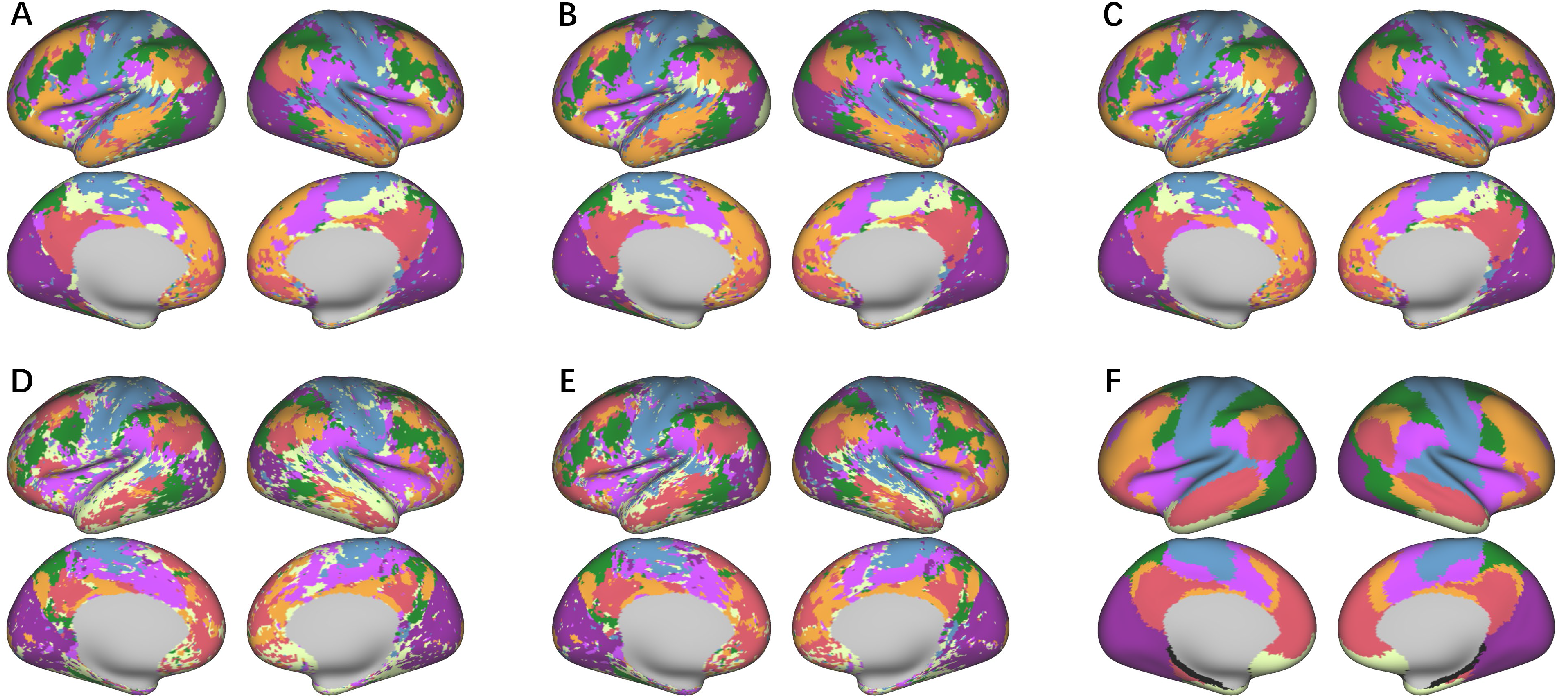
The brain network partitioning results for the representative subject. A: SDAE pre-training + *k*-means clustering. B: SDAE pretraining + DEC clustering. C: SDAE pretraining + IDEC clustering. D: MVM pretraining + *k*-means clustering. E: Indiv-Brain. F: yeo7 template[1].

The major improvement in our model compared to the traditional DEC model is the introduction of the training tasks concept from the BERT model and integrating the attention mechanism from the transformer encoder. This allows each node to reference and utilize information from other nodes during representation learning, aligning with the biological properties of the brain where different nodes are interrelated and influence each other. This integration of node information is primarily reflected through the attention mechanism.

### 5.3 In-depth Analysis of Attention Scores

The Fig. 4 illustrates the attention visualization for the representative subject using optimal parameters after training. Fig. 4A shows the first attention block, and Fig. 4B shows the second attention block. After MVM pretraining, the attention matrix (i.e., the term softmax 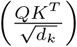 in Eq. (2)) for the selected tokens has values only in a few columns, as the others do not participate in the loss calculation. In these few columns, gaps appear in the rows corresponding to the selected tokens, which are used to compute the reconstructed tokens. In the V matrix of the attention mechanism, the selected masked tokens contain no information due to the masking process. The resulting matrix, after multiplying these two matrices, shows that unselected tokens do not contain meaningful information as they do not participate in the loss calculation, while the selected tokens successfully reconstruct their signals using information from other tokens. In Block 2, apart from the decreased diagonal values, the basic pattern is similar to that of Block 1, indicating that our model achieves the intended goal of constructing self-representation using the relationship with other tokens.

**Figure 4:**
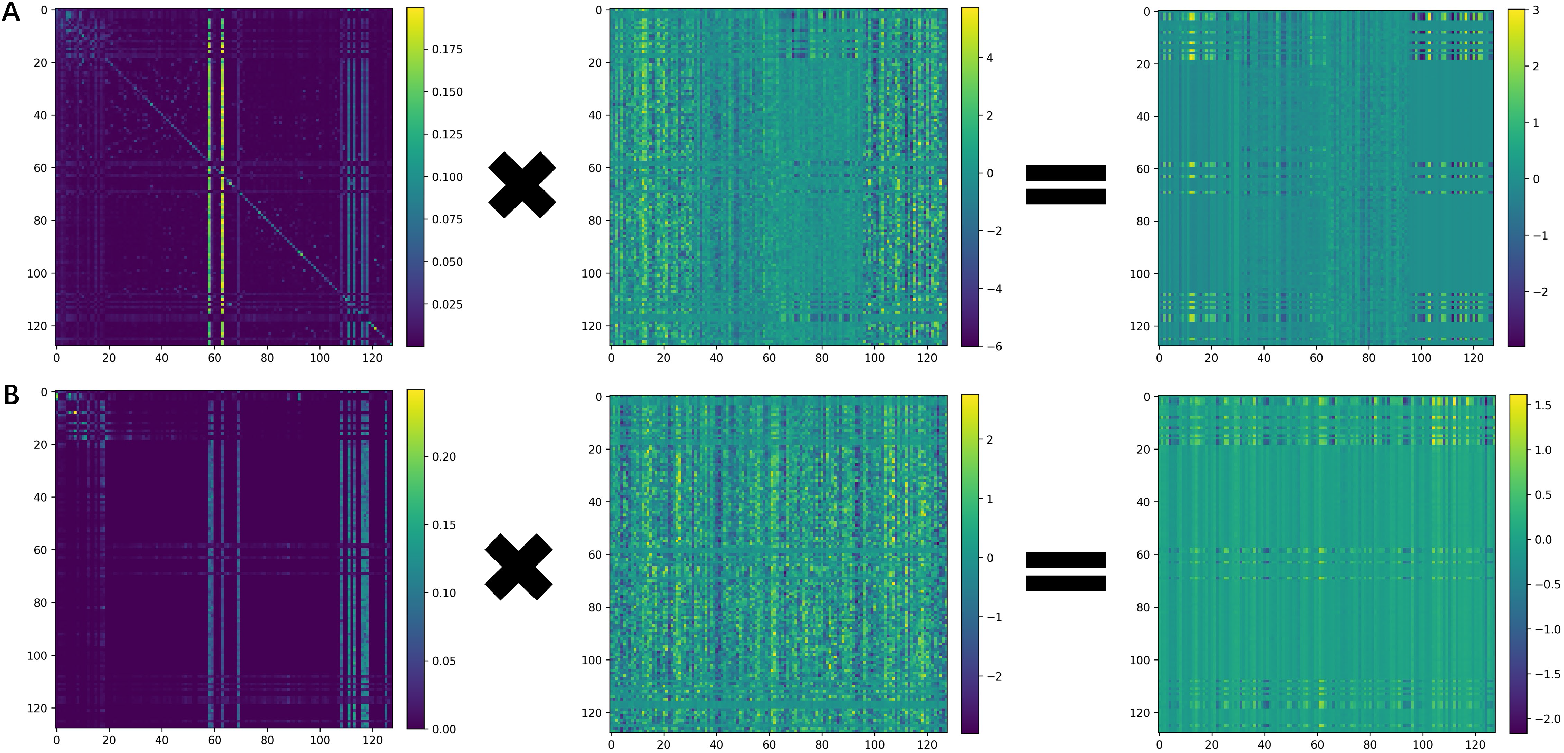
Attention visualization. We visualized the operation process of the attention blocks in the transformer encoder for the optimal hyperparameter combination results of the representative subject. The three figures of each row are softmax 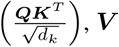, and the output of the attention block. A: the first attention block; B: the second attention block.

## 6 Discussion and Conclusion

### Discussion

The Indiv-Brain model proposed in this study achieves individualized brain network partitioning by applying deep learning to individual fMRI data. Compared to other methods, Indiv-Brain demonstrates significant advantages in several aspects. First, this study overcomes the limitations of conventional brain region partitioning methods by avoiding the use of predefined brain parcellations. Traditional methods rely on group data for model training, ignoring individual differences, which may result in poor adaptability to special subjects. Indiv-Brain, by directly using individual fMRI data for training, ensures precise partitioning of individual brain networks. Secondly, the study introduces a Transformer-based attention mechanism into the model structure. This innovation enables the model to capture the interactions between nodes, reflecting the complex interactions between different brain regions. In the process of sentence construction, the use of correlation coefficients allows consideration of long-distance related nodes. The introduction of the attention mechanism significantly enhances the model’s representational capacity, making the features of each node more biologically meaningful and interpretable. In terms of performance evaluation, experimental results show that Indiv-Brain comprehensively surpasses traditional DEC models and their improved versions IDEC in accuracy. This not only validates the effectiveness of our model in individualized brain network partitioning but also indicates the potential of combining deep embedding clustering with attention mechanisms to improve clustering accuracy. The Indiv-Brain model provides an effective method for individualized brain network partitioning, paving new paths for neuroscience research and clinical applications. We believe that with the development of deep learning technologies and the conduct of more research, individualized brain network studies will offer deeper insights into the complex functions and individual differences of the human brain.

### Conclusion

In summary, the proposed Indiv-Brain model has made significant progress in the field of individualized brain network partitioning. By combining SDAE, MVM, and DEC-A, this model efficiently achieves precise partitioning of individual brain networks without relying on predefined brain parcellations. Experimental results show that Indiv-Brain not only significantly outperforms traditional methods in accuracy but also possesses advantages in model interpretability and adaptability.

Future work will focus on further optimizing the model structure and algorithms to improve its adaptability and robustness across different subjects and pathological data. Additionally, we will explore the application of the model to other neuroimaging data, such as MEG, to provide more comprehensive support for understanding individual brain functional networks.

### Limitation

Due to time and computing resource constraints, we were unable to perform a comprehensive hyperparameter search experiments for subjects other than those included in our study. Additionally, the number of subjects used to train our model was relatively small, which limits the persuasiveness of our model. Furthermore, in terms of clustering methods, we only employed DEC clustering based on KL divergence. In future work, we can explore other deep clustering methods, such as spectral clustering or mutual information-based clustering. Lastly, we can experiment with the Yeo17 template to further investigate the robustness of our model.

### Broader Impacts

We believe that individualized brain network partitioning has promising applications in individual difference research as large models develop, and it will be a non-negligible component of the study to understand our cognitive process. However, governmental regulations and efforts from research communities are required to ensure the privacy of one’s biological data.

## Appendix

### A Data preprocessing

#### Human connectome project

In this study, we utilized resting state functional magnetic resonance imaging (rs-fMRI) data from the 1200 Subjects Release (S1200) dataset within the Human Connectome Project [20] (HCP). For each subject, rs-fMRI data were acquired in two sessions, with two runs separately acquired per session. Within each session, the phase encoding directions of the two runs are opposite: left-to-right (LR) and right-to-left (RL). Each run was acquired with a gradient-echo planar imaging sequence with following parameters: time repetition (TR) = 720 *ms*; time echo (TE) = 33.1 *ms*; flip angle = 52^°^; field of view=208×180 *mm*^2^; matrix=104×90; voxel size=2×2×2 *mm*^2^; multiband factor=8 and 1200 volumes (i.e., 14.4 min). For more detailed information about the data acquisition protocols, refer to [20]. Written informed consent was obtained from all participants, and the scanning protocol was approved by the Institutional Review Board of Washington University. The dataset containing de-identified data is publicly available on the ConnectomeDB database (https://db.humanconnectome.org). For our experiments, we selected 8 individuals from the “100 Unrelated Subjects” group within the S1200 release without family structure issues. We used data from one run (session 1, LR phase encoding) for each subject.

##### Preprocessing

All rs-fMRI data were preprocessed using HCP minimal preprocessing pipeline [43]. Briefly, all images data underwent gradient distortion correction, motion correction, echo planar image (EPI) distortion correction, registration to the Montreal Neurological Institute (MNI) space from the original functional image in one spline resampling step and intensity normalization. Then, the volume time series were mapped to the standard CIFTI grayordinate space, resampled from high-resolution mesh to the downsampled 32k_fs_LR mesh and smoothed using a 2 mm full width at half maximum (FWHM) smoothing kernel on the surface. The result CIFTI file for each subject followed the file naming pattern: {Subject_ID}_REST1_LR _Atlas_MSMAll.dtseries.nii. Moreover, we applied the following additional preprocessing steps: i) linear trend removal, ii) confound removal by regressing out 36 parameters including 9 parameters (6 motion parameters *x, y, z* translations and rotations), mean global, white matter (WM) and cerebrospinal fluid (CSF) time series), temporal derivative of 9 parameters across the time series and the quadratic term of the 18 parameters, iii) temporal band-pass filtering (0.01–0.1 Hz), iv) scrubbing. Frames with framewise displacement exceeding 0.5 *mm* and their adjacent volumes (1 back and 2 forward) were replaced with linearly interpolated data. Subject with more than 25% interpolated volumes was excluded. Finally, we normalized the time series to have a mean of 0 and unit variance. After all the above steps, no subject was excluded.

**Figure 5:**
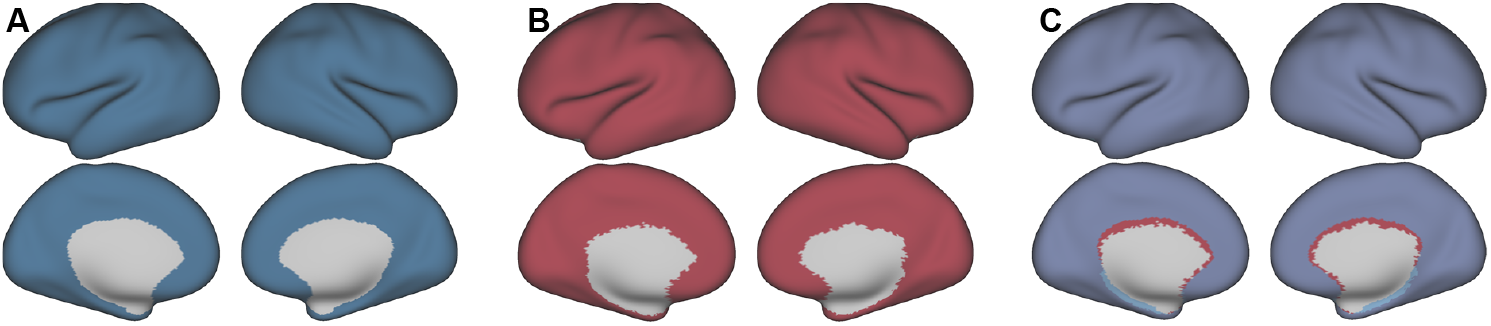
**Left:**. The standard fs_LR_32k template. **Middle:** The Yeo template from [38]. **Right:**.The overlap of the two prior templates.

### B Other subjects

**Figure 6:**
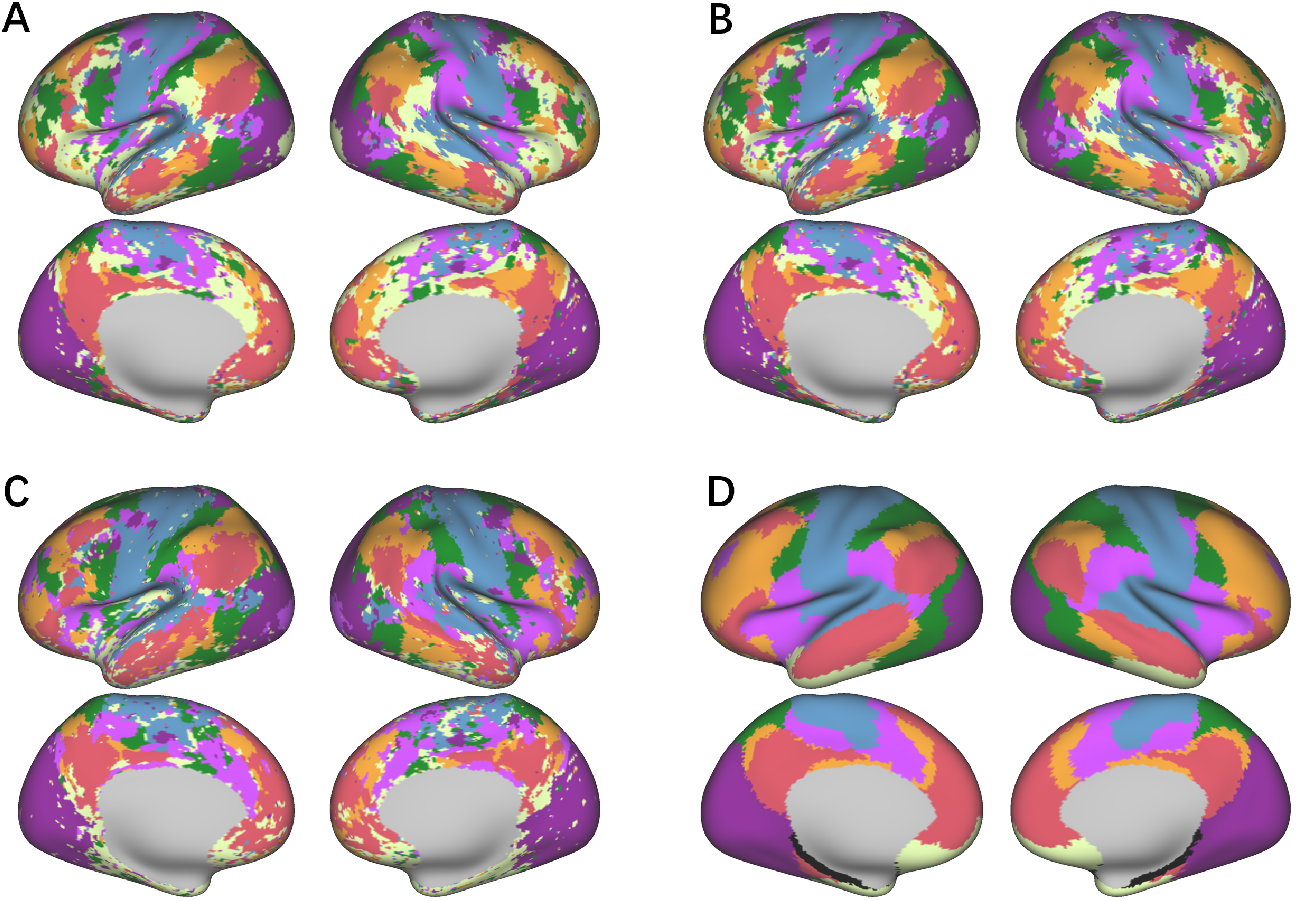
The brain network partitioning results for subject 2. A: SDAE pretraining + DEC clustering. B: SDAE pretraining + IDEC clustering. C: Indiv-Brain. D: yeo7 template [38].

**Figure 7:**
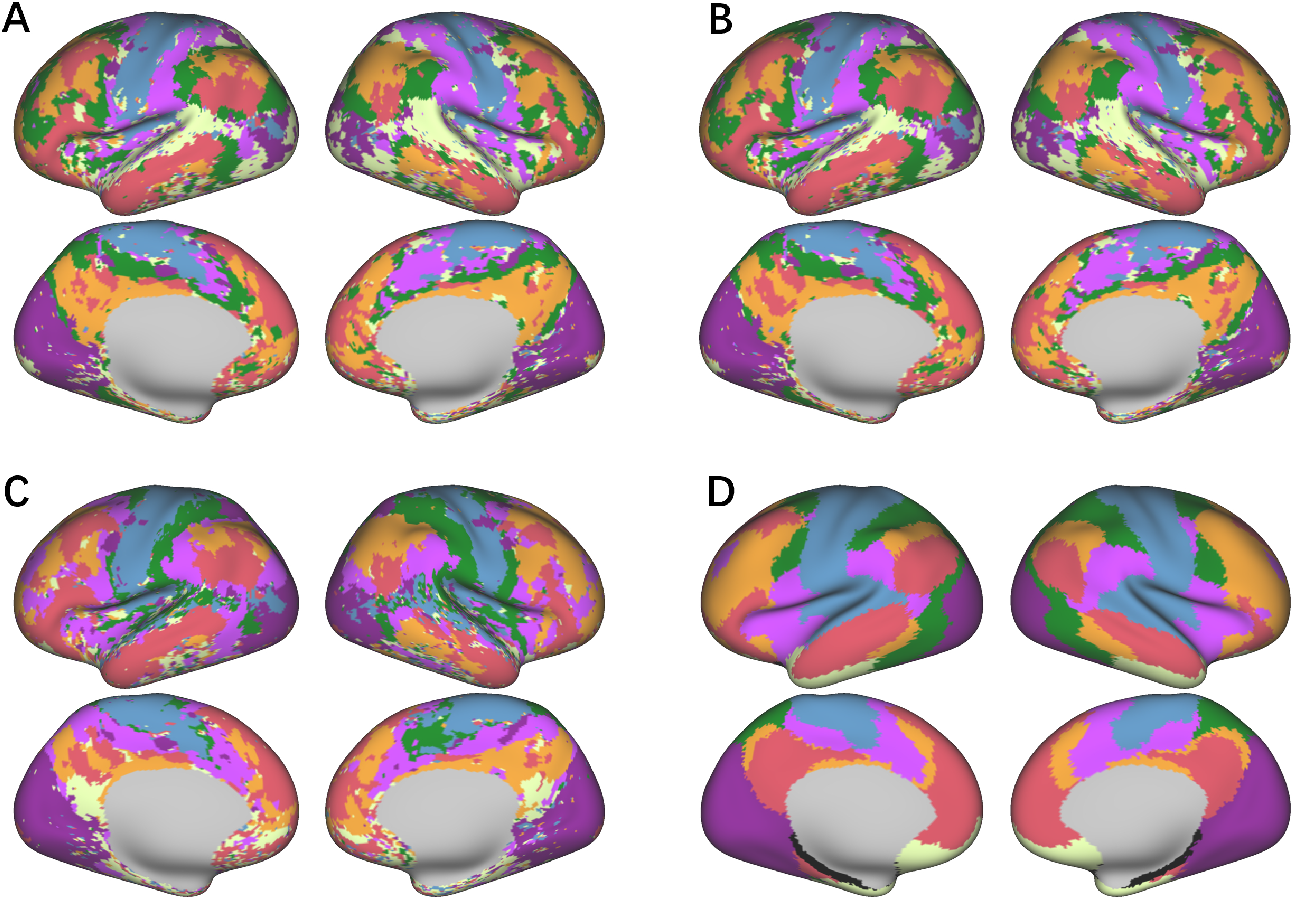
The brain network partitioning results for subject 3. A: SDAE pretraining + DEC clustering. B: SDAE pretraining + IDEC clustering. C: Indiv-Brain. D: yeo7 template [38].

**Figure 8:**
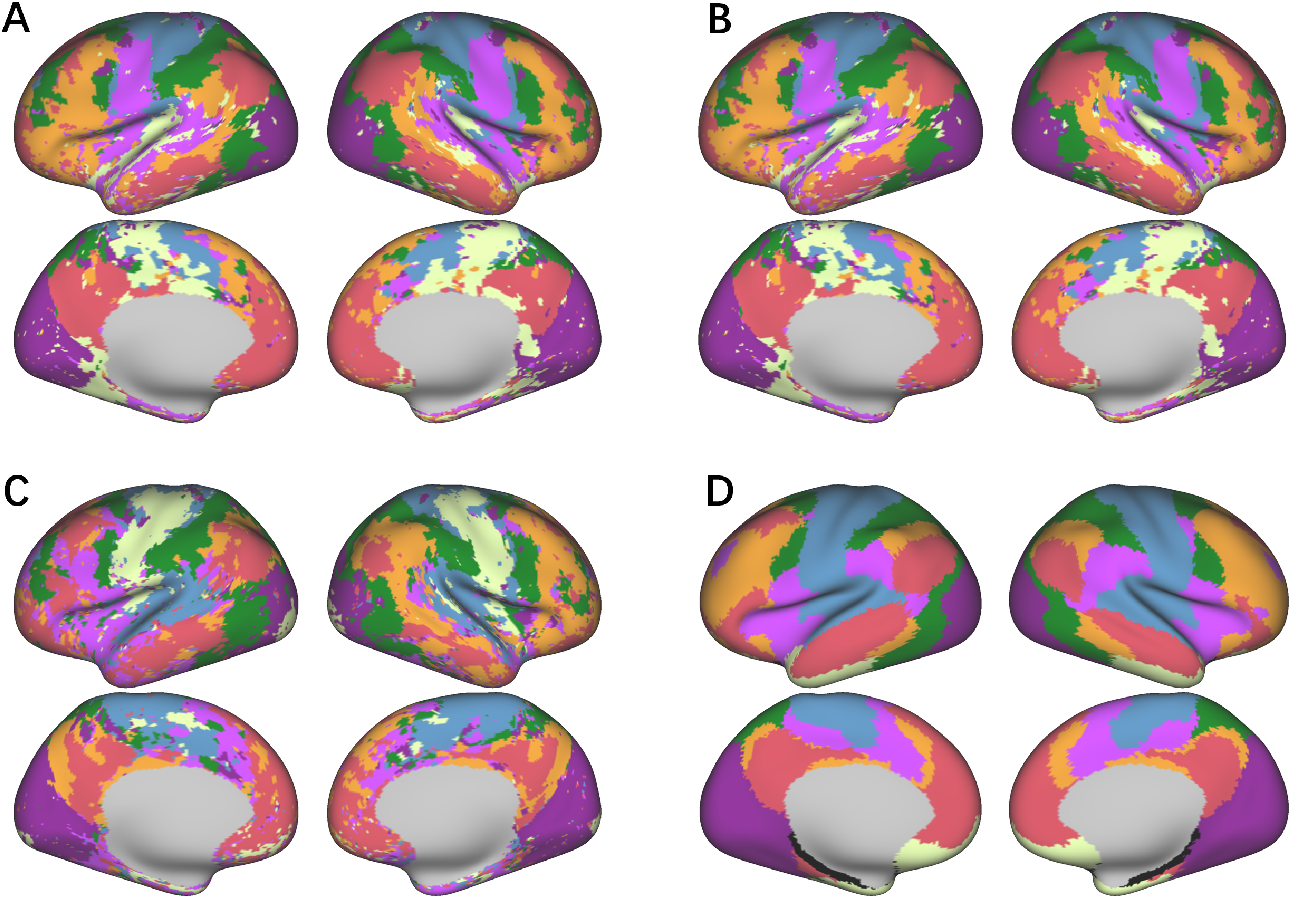
The brain network partitioning results for subject 4. A: SDAE pretraining + DEC clustering. B: SDAE pretraining + IDEC clustering. C: Indiv-Brain. D: yeo7 template [38].

**Figure 9:**
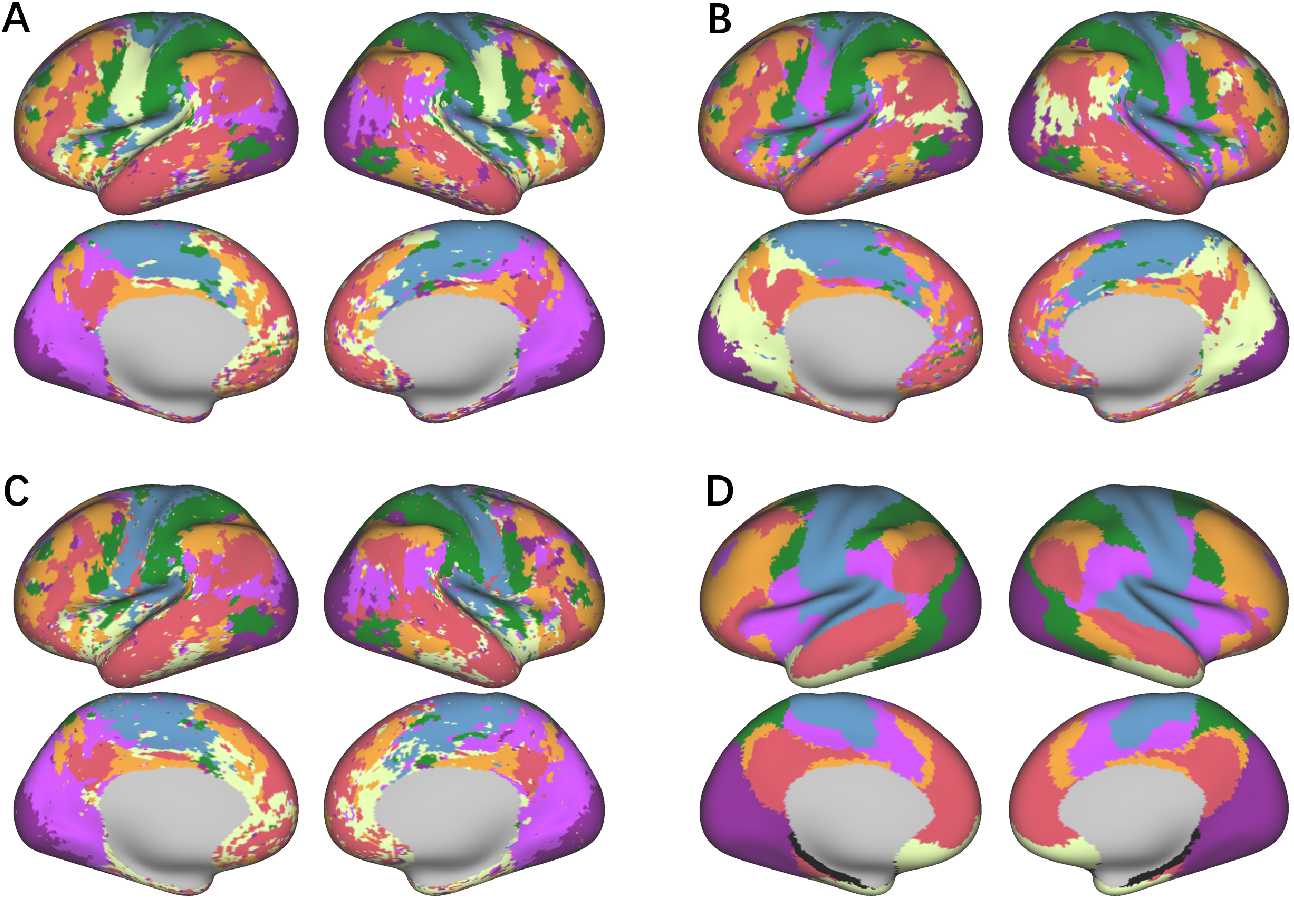
The brain network partitioning results for subject 5. A: SDAE pretraining + DEC clustering. B: SDAE pretraining + IDEC clustering. C: Indiv-Brain. D: yeo7 template [38].

**Figure 10:**
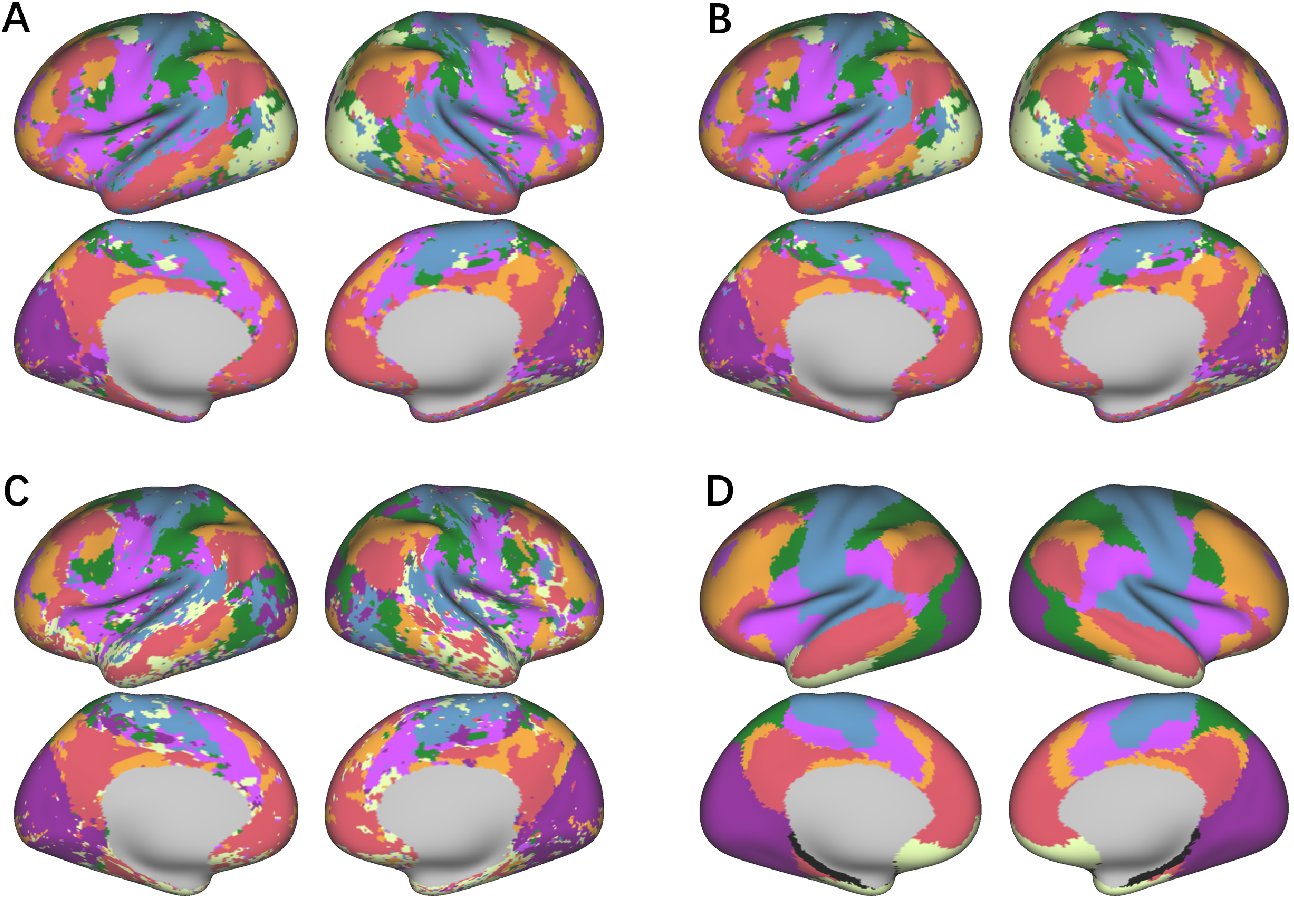
The brain network partitioning results for subject 6. A: SDAE pretraining + DEC clustering. B: SDAE pretraining + IDEC clustering. C: Indiv-Brain. D: yeo7 template [38].

**Figure 11:**
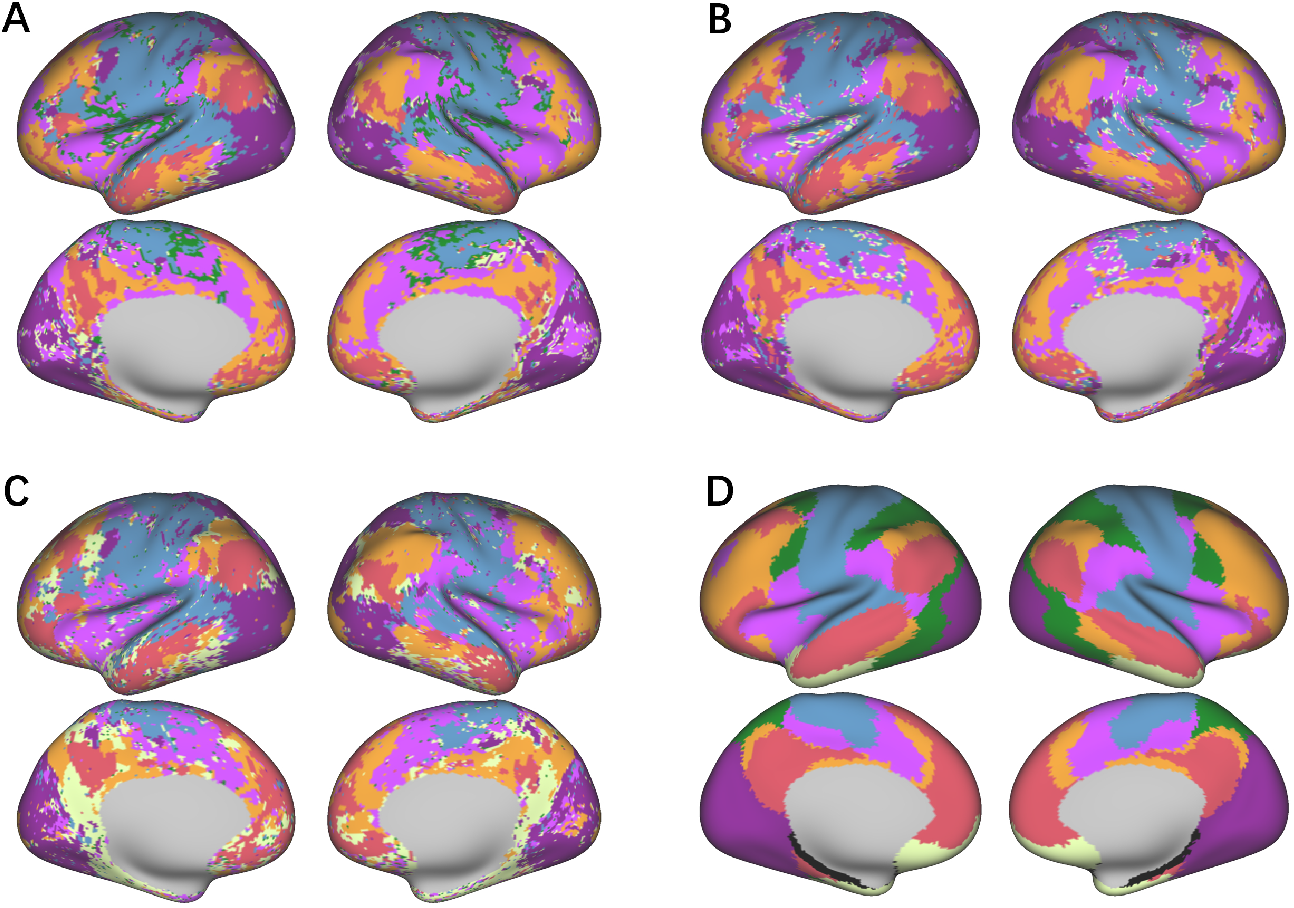
The brain network partitioning results for subject 7. A: SDAE pretraining + DEC clustering. B: SDAE pretraining + IDEC clustering. C: Indiv-Brain. D: yeo7 template [38].

**Figure 12:**
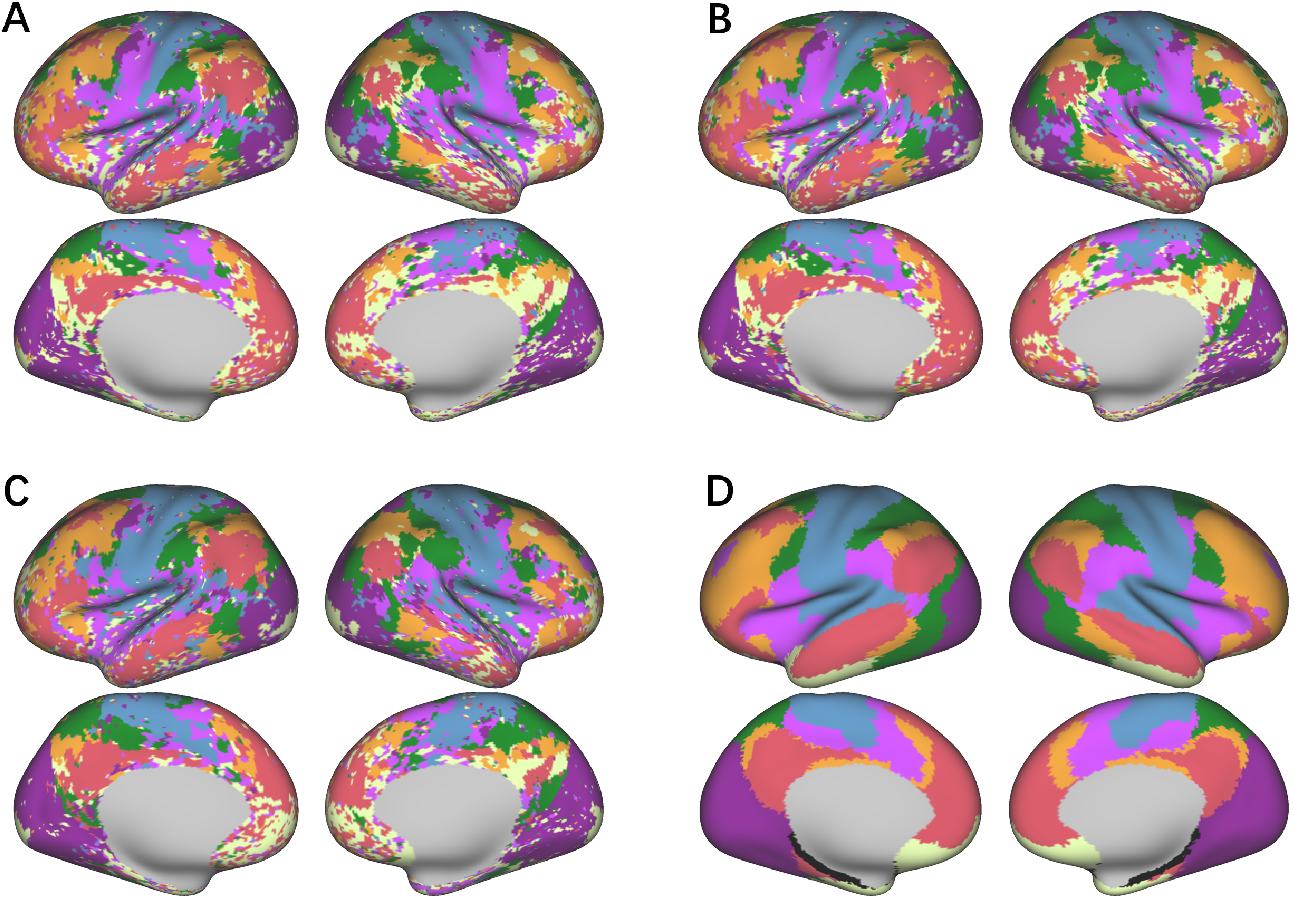
The brain network partitioning results for subject 8. A: SDAE pretraining + DEC clustering. B: SDAE pretraining + IDEC clustering. C: Indiv-Brain. D: yeo7 template [38].

## Notes

https://db.humanconnectome.org

### Competing Interest Statement

The authors have declared no competing interest.

## References

1 B. T. Thomas Yeo, Fenna M. Krienen, Jorge Sepulcre, Mert R. Sabuncu, Danial Lashkari, Marisa Hollinshead, Joshua L. Roffman, Jordan W. Smoller, Lilla Zöllei, Jonathan R. Polimeni, Bruce Fischl, Hesheng Liu, and Randy L. Buckner. The organization of the human cerebral cortex estimated by intrinsic functional connectivity. Journal of Neurophysiology, 106(3):1125–1165, 2011.

2 Jonathan D. Power, Alexander Li Cohen, Steven M. Nelson, Gagan S. Wig, Kelly Anne Barnes, Jessica A. Church, Alecia C. Vogel, Timothy O. Laumann, Francis M. Miezin, Bradley L. Schlaggar, and Steven E. Petersen. Functional network organization of the human brain. Neuron, 72:665–678, 2011.

3 Evan M. Gordon, Timothy O. Laumann, Babatunde Adeyemo, Jeremy F. Huckins, William M. Kelley, and Steven E. Petersen. Generation and Evaluation of a Cortical Area Parcellation from Resting-State Correlations. Cerebral Cortex, 26(1):288–303, 2014.

4 Nico U. F. Dosenbach, Damien A. Fair, Francis M. Miezin, Alexander L. Cohen, Kristin K. Wenger, Ronny A. T. Dosenbach, Michael D. Fox, Abraham Z. Snyder, Justin L. Vincent, Marcus E. Raichle, Bradley L. Schlaggar, and Steven E. Petersen. Distinct brain networks for adaptive and stable task control in humans. Proceedings of the National Academy of Sciences, 104(26):11073–11078, 2007.

5 Gaëlle Doucet, Mikaël Naveau, Laurent Petit, Nicolas Delcroix, Laure Zago, Fabrice Crivello, Gaël Jobard, Nathalie Tzourio-Mazoyer, Bernard Mazoyer, Emmanuel Mellet, and Marc Joliot. Brain activity at rest: a multiscale hierarchical functional organization. Journal of Neurophysiology, 105(6):2753–2763, 2011.

6 Gagan S. Wig. Segregated systems of human brain networks. Trends in Cognitive Sciences, 21 (12):981–996, 2017.

7 Olaf Sporns. Network attributes for segregation and integration in the human brain. Current Opinion in Neurobiology, 23(2):162–171, 2013.

8 Janine Diane Bijsterbosch, Mark W Woolrich, Matthew F Glasser, Emma C Robinson, Christian F Beckmann, David C Van Essen, Samuel J Harrison, and Stephen M Smith. The relationship between spatial configuration and functional connectivity of brain regions. eLife, 7:e32992, 2018.

9 Evan M. Gordon, Timothy O. Laumann, Adrian W. Gilmore, Dillan J. Newbold, Deanna J. Greene, Jeffrey J. Berg, Mario Ortega, Catherine Hoyt-Drazen, Caterina Gratton, Haoxin Sun, Jacqueline M. Hampton, Rebecca S. Coalson, Annie L. Nguyen, Kathleen B. McDermott, Joshua S. Shimony, Abraham Z. Snyder, Bradley L. Schlaggar, Steven E. Petersen, Steven M. Nelson, and Nico U. F. Dosenbach. Precision Functional Mapping of Individual Human Brains. Neuron, 95(4):791–807.e7, 2017.

10 Xuhong Liao, Miao Cao, Mingrui Xia, and Yong He. Individual differences and time-varying features of modular brain architecture. NeuroImage, 152:94–107, 2017.

11 Zhitao Guo, Xiaojie Zhao, Li Yao, and Zhiying Long. Improved brain community structure detection by two-step weighted modularity maximization. PLOS ONE, 18(12):e0295428, 2023.

12 Qawi K. Telesford, Mary-Ellen Lynall, Jean Vettel, Michael B. Miller, Scott T. Grafton, and Danielle S. Bassett. Detection of functional brain network reconfiguration during task-driven cognitive states. NeuroImage, 142:198–210, 2016.

13 U Braun, A Schafer, H Walter, S Erk, N Romanczuk-Seiferth, L Haddad, JI Schweiger, O Grimm, A Heinz, H Tost, A Meyer-Lindenberg, and DS Bassett. Dynamic reconfiguration of frontal brain networks during executive cognition in humans. Proc Natl Acad Sci U S A, 112 (37):11678–11683, 2015.

14 Hongming Li, Dhivya Srinivasan, Chuanjun Zhuo, Zaixu Cui, Raquel E. Gur, Ruben C. Gur, Desmond J. Oathes, Christos Davatzikos, Theodore D. Satterthwaite, and Yong Fan. Computing personalized brain functional networks from fmri using self-supervised deep learning. Medical Image Analysis, 85:102756, 2023.

15 Ru Kong, Jingwei Li, Csaba Orban, Mert R Sabuncu, Hesheng Liu, Alexander Schaefer, Nanbo Sun, Xi-Nian Zuo, Avram J Holmes, Simon B Eickhoff, and BT Thomas Yeo. Spatial Topography of Individual-Specific Cortical Networks Predicts Human Cognition, Personality, and Emotion. Cerebral Cortex, 29(6):2533–2551, 2018.

16 Hongming Li, Theodore D. Satterthwaite, and Yong Fan. Large-scale sparse functional networks from resting state fmri. NeuroImage, 156:1–13, 2017.

17 Samuel J. Harrison, Mark W. Woolrich, Emma C. Robinson, Matthew F. Glasser, Christian F. Beckmann, Mark Jenkinson, and Stephen M. Smith. Large-scale probabilistic functional modes from resting state fmri. NeuroImage, 109:217–231, 2015.

18 Samuel J. Harrison, Janine D. Bijsterbosch, Andrew R. Segerdahl, Sean P. Fitzgibbon, Seyedeh-Rezvan Farahibozorg, Eugene P. Duff, Stephen M. Smith, and Mark W. Woolrich. Modelling subject variability in the spatial and temporal characteristics of functional modes. NeuroImage, 222:117226, 2020.

19 Hongming Li, Xiaofeng Zhu, and Yong Fan. Identification of multi-scale hierarchical brain functional networks using deep matrix factorization. In Medical Image Computing and Computer Assisted Intervention – MICCAI 2018, pages 223–231, 2018.

20 David C. Van Essen, Stephen M. Smith, Deanna M. Barch, Timothy E.J. Behrens, Essa Yacoub, and Kamil Ugurbil. The wu-minn human connectome project: An overview. NeuroImage, 80: 62–79, 2013.

21 Junyuan Xie, Ross Girshick, and Ali Farhadi. Unsupervised deep embedding for clustering analysis. In Proceedings of the 33rd International Conference on International Conference on Machine Learning - Volume 48, page 478–487, 2016.

22 Bo Yang, Xiao Fu, Nicholas D. Sidiropoulos, and Mingyi Hong. Towards k-means-friendly spaces: simultaneous deep learning and clustering. In Proceedings of the 34th International Conference on Machine Learning, page 3861–3870, 2017.

23 Zhuxi Jiang, Yin Zheng, Huachun Tan, Bangsheng Tang, and Hanning Zhou. Variational deep embedding: an unsupervised and generative approach to clustering. In Proceedings of the 26th International Joint Conference on Artificial Intelligence, page 1965–1972, 2017.

24 Mathilde Caron, Piotr Bojanowski, Armand Joulin, and Matthijs Douze. Deep clustering for unsupervised learning of visual features. In Proceedings of the European Conference on Computer Vision (ECCV), 2018.

25 Pan Ji, Tong Zhang, Hongdong Li, Mathieu Salzmann, and Ian Reid. Deep subspace clustering networks. In Proceedings of the 31st International Conference on Neural Information Processing Systems, page 23–32, 2017.

26 Jacob Devlin, Ming-Wei Chang, Kenton Lee, and Kristina Toutanova. Bert: Pre-training of deep bidirectional transformers for language understanding. In North American Chapter of the Association for Computational Linguistics, 2019.

27 Kaiming He, Xinlei Chen, Saining Xie, Yanghao Li, Piotr Doll’ar, and Ross B. Girshick. Masked autoencoders are scalable vision learners. 2022 IEEE/CVF Conference on Computer Vision and Pattern Recognition (CVPR), pages 15979–15988, 2021.

28 Zhenda Xie, Zheng Zhang, Yue Cao, Yutong Lin, Jianmin Bao, Zhuliang Yao, Qi Dai, and Han Hu. Simmim: a simple framework for masked image modeling. In 2022 IEEE/CVF Conference on Computer Vision and Pattern Recognition (CVPR), pages 9643–9653, 2022.

29 Shuangfei Zhai, Navdeep Jaitly, Jason Ramapuram, Dan Busbridge, Tatiana Likhomanenko, Joseph Y. Cheng, Walter Talbott, Chen Huang, Hanlin Goh, and Joshua M. Susskind. Position Prediction as an Effective Pretraining Strategy. In Proceedings of the 39th International Conference on Machine Learning, pages 26010–26027, 2022.

30 Geoffrey E. Hinton and Richard S. Zemel. Autoencoders, minimum description length and helmholtz free energy. In Proceedings of the 6th International Conference on Neural Information Processing Systems, page 3–10, 1993.

31 Pascal Vincent, Hugo Larochelle, Isabelle Lajoie, Yoshua Bengio, and Pierre-Antoine Manzagol. Stacked denoising autoencoders: Learning useful representations in a deep network with a local denoising criterion. J. Mach. Learn. Res., 11:3371–3408, 2010.

32 Hangbo Bao, Li Dong, Songhao Piao, and Furu Wei. BEiT: BERT pre-training of image transformers. In International Conference on Learning Representations, 2022.

33 Armin W. Thomas, Christopher Ré, and Russell A. Poldrack. Self-Supervised Learning of Brain Dynamics from Broad Neuroimaging Data. In Advances in Neural Information Processing Systems, 2022.

34 Zijiao Chen, Jiaxin Qing, Tiange Xiang, Wan Lin Yue, and Juan Helen Zhou. Seeing Beyond the Brain: Conditional Diffusion Model with Sparse Masked Modeling for Vision Decoding. In 2023 IEEE/CVF Conference on Computer Vision and Pattern Recognition (CVPR), pages 22710–22720, 2023.

35 Xifeng Guo, Long Gao, Xinwang Liu, and Jianping Yin. Improved deep embedded clustering with local structure preservation. In Proceedings of the 26th International Joint Conference on Artificial Intelligence, page 1753–1759, 2017.

36 Xuan-Bac Nguyen, Duc Toan Bui, Chi Nhan Duong, Tien D. Bui, and Khoa Luu. Clusformer: A transformer based clustering approach to unsupervised large-scale face and visual landmark recognition. In 2021 IEEE/CVF Conference on Computer Vision and Pattern Recognition (CVPR), pages 10842–10851, 2021.

37 Ashish Vaswani, Noam Shazeer, Niki Parmar, Jakob Uszkoreit, Llion Jones, Aidan N. Gomez, Łukasz Kaiser, and Illia Polosukhin. Attention is all you need. In Proceedings of the 31st International Conference on Neural Information Processing Systems, page 6000–6010, 2017.

38 David C. Van Essen, John Smith, Matthew F. Glasser, Jennifer Elam, Chad J. Donahue, Donna L. Dierker, Erin K. Reid, Timothy Coalson, and John Harwell. The brain analysis library of spatial maps and atlases (balsa) database. NeuroImage, 144:270–274, 2017.

39 Iulia Turc, Ming-Wei Chang, Kenton Lee, and Kristina Toutanova. Well-read students learn better: On the importance of pre-training compact models. arXiv preprint 1908.08962, 2019.

40 Dan Hendrycks and Kevin Gimpel. Gaussian error linear units (gelus). arXiv preprint 1606.08415, 2016.

41 Adam Paszke, Sam Gross, Francisco Massa, Adam Lerer, James Bradbury, Gregory Chanan, Trevor Killeen, Zeming Lin, Natalia Gimelshein, Luca Antiga, Alban Desmaison, Andreas Köpf, Edward Yang, Zach DeVito, Martin Raison, Alykhan Tejani, Sasank Chilamkurthy, Benoit Steiner, Lu Fang, Junjie Bai, and Soumith Chintala. Pytorch: an imperative style, high-performance deep learning library. In Proceedings of the 33rd International Conference on Neural Information Processing Systems, 2019.

42 Y. Lecun, L. Bottou, Y. Bengio, and P. Haffner. Gradient-based learning applied to document recognition. Proceedings of the IEEE, 86(11):2278–2324, 1998.

43 Matthew F. Glasser, Stamatios N. Sotiropoulos, J. Anthony Wilson, Timothy S. Coalson, Bruce Fischl, Jesper L. Andersson, Junqian Xu, Saad Jbabdi, Matthew Webster, Jonathan R. Polimeni, David C. Van Essen, and Mark Jenkinson. The minimal preprocessing pipelines for the human connectome project. NeuroImage, 80:105–124, 2013.

